# Imaging Multiple Sclerosis Pathology at 160μm Isotropic Resolution by Human Whole-Brain *Ex Vivo* Magnetic Resonance Imaging at 3T

**DOI:** 10.1101/2021.01.03.425097

**Authors:** Weigel Matthias, Dechent Peter, Galbusera Riccardo, Bahn Erik, Nair Govind, Kappos Ludwig, Brück Wolfgang, Stadelmann Christine, Granziera Cristina

## Abstract

Postmortem magnetic resonance imaging (MRI) of the fixed healthy and diseased human brain facilitates spatial resolutions and image quality that is not achievable with *in vivo* MRI scans. Though challenging - and almost exclusively performed at 7T field strength - depicting the tissue architecture of the entire brain in fine detail is invaluable since it enables the study of neuroanatomy and uncovers important pathological features in neurological disorders. The objectives of the present work were (i) to develop a 3D isotropic ultra-high-resolution imaging approach for human whole-brain *ex vivo* acquisitions working on a standard clinical 3T MRI system, and (ii) to explore the sensitivity and specificity of this concept for specific pathoanatomical features of multiple sclerosis. The reconstructed images demonstrate unprecedented resolution and soft tissue contrast of the diseased human brain at 3T, thus allowing visualization of sub-millimetric lesions in the different cortical layers and in the cerebellar cortex, as well as unique cortical lesion characteristics such as the presence of incomplete / complete iron rims, and patterns of iron accumulation. Further details such as the subpial molecular layer, the line of Gennari, and some intrathalamic nuclei are also well distinguishable.

## Introduction

Postmortem magnetic resonance (MR) imaging of the formalin-fixed healthy and diseased human brain offers the possibility to gain a deep understanding of neuromorphology and neuropathology, including the option to compare macro- and microanatomical and tissue-related features to histopathology ^1–3^. Indeed, postmortem imaging benefits from a number of advantages compared to *in vivo* imaging such as the lack of many sources of artifacts and the opportunity to perform long scan sessions (i.e., ranging from hours to days), which enable spatial resolutions and image quality not achievable with *in vivo* scans ^2,4–6^. Postmortem imaging of fixed brains, however, is also characterized by several challenges, encompassing the changes in MRI properties due to fixation, large image artifacts resulting from the magnetic susceptibility differences that occur at (residual) air - tissue boundaries, etc. ^1,7–9^.

Yet, postmortem whole brain imaging is important as it is required to observe neuroanatomic relationships across distant brain regions, to provide neuroanatomical reference points in standard stereotactic space and to understand neurological diseases affecting the entire brain structure.

In contrast to the imaging of tissue-blocks or slices ^10–12^ – which is usually performed in small-bore scanners and/or using specialized receiver-coils – *ex vivo* whole-brain MR imaging has been frequently performed at 7T main field strength ^4,13^, since it generally provides a higher signal-to-noise-ratio (SNR) per unit time. SNR is usually the most limiting factor in terms of achievable resolution; and increasing SNR solely by signal averaging leads to an unfavorable quadratic increase of acquisition time. In contrast, 3T MR systems have the benefit of being much more widely available, are more economic in operation, provide quite homogeneous excitation and reception fields and, not least, the acquired *ex vivo* image data allow more direct comparisons with patient data from clinical routine.

The purpose of this work was to develop – and to test for the viable boundaries of – an ultra-high-resolution MR acquisition for human whole-brain *ex vivo* imaging with strong soft tissue contrast on a standard clinical 3T MR system. For this purpose, we studied the entire brain of a 58-year-old patient who was affected by multiple sclerosis for 24-years and explored the sensitivity and specificity of the acquired ultra-high-resolution images to specific pathological features of this common disease, both in the white matter and in the cortical brain tissue. Four different base protocols were established to acquire isotropic 3D resolutions between 160μm and 240μm within a few hours up to a few days. The resulting MR images display unprecedented quality of the diseased human MS brain at 3T field strength.

## Materials and Methods

### Specimen preparation and experimental setup

The following experiments were approved by the ethical review committee of the Medical Center of Göttingen (29/9/10). Clinical data and brain tissue were collected following established standard operating procedures (SOPs) developed within the KKNMS (German Competence network for multiple sclerosis).

The whole brain of a 58-year-old man with a 24-year history of multiple sclerosis was investigated. The autopsy had been performed 24 hours postmortem and the brain was fixated directly in 4% neutral buffered formaldehyde solution (formalin). Afterwards, the brain was transferred to a custom-built, dome-shaped container as depicted in Refs. ^3,14,15^. Within the MRI-compatible container the brain was immersed in Fomblin^®^ perfluoropolyether (Solvay Specialty Polymers USA, LLC, West Deptford, NJ, USA). This fluoropolymer has a magnetic susceptibility similar to that of tissue and lacks any signal in hydrogen-based MRI ^16^. Since air bubbles produce substantial susceptibility artifacts, these were removed through the degassing ports of the container using a vacuum pump ^3,14,15^.

All experiments were performed on a 3T whole-body MR system (Prisma^Fit^, Siemens Healthineers, Erlangen, Germany) with a maximal gradient amplitude of 80mT/m and a maximal slew rate of 200mT/m/ms. For radiofrequency (RF) transmission the built-in body coil was employed. For RF reception the standard, manufacturer-supplied 20-channel phased-array head and neck coil was used.

The overall protocol consisted of initial localizer scans for controlling the position and alignment of the brain and its container both in the RF head and neck coil and in the scanner isocenter. Subsequently, a series of acquisitions with one of the here suggested ultra-high-resolution MR imaging (URI) acquisition protocols was performed. The maximum number of protocol repetitions was defined by the whole time that was available in our respective measurement slot. The shortest measurement slot had ~56h, the longest had ~90h (cf. Supplementary Tab. S1).

### The 3T URI acquisition approach

Four URI base protocols with isotropic 3D resolutions 160μm, 180μm, 200μm, and 240μm were established. The protocols are based on the principle of a common RF spoiled gradient echo sequence, often referred to as a fast low-angle shot (FLASH) sequence ^17^. Since vendor MRI sequences are typically restricted regarding what protocol parameters can be modified and which parameter range is permitted, an in-house developed FLASH sequence with a simple linear 3D phase encoding loop structure was employed. As an additional benefit, with the in-house sequence a more precise monitoring of the needed MR system performance was feasible.

All four base protocols used a constant echo time TE = 18ms, a constant receiver bandwidth of 50 Hz/Px, and transversal slice orientation (no angulation). This overall procedure ensured images of similar contrast and imaging behavior. The resolution-dependent parameters were: (1) Isotropic 3D resolution (160μm)^3^, field-of-view (FOV) = 192×150cm^2^, matrix = 1200×936, slice thickness = 0.16mm, slices per slab = 704, phase encoding direction right to left, flip angle = 31deg, TR = 38ms, TA_base_ = 06:57:20h; (2) Isotropic 3D resolution (180μm)^3^, FOV = 192×161cm^2^, matrix = 1072×896, slice thickness = 0.18mm, slices per slab = 768, phase encoding direction right to left, slice oversampling = 80%, flip angle = 31deg, TR = 37ms, TA_base_ = 07:04:49h; (3) Isotropic 3D resolution (200μm)^3^, FOV = 192×156cm^2^, matrix = 960×780, slice thickness = 0.20mm, slices per slab = 768, phase encoding direction right to left, slice oversampling = 33.3%, flip angle = 31deg, TR = 37ms, TA_base_ = 08:12:33h; (4) Isotropic 3D resolution (240μm)^3^, FOV = 192×190cm^2^, matrix = 800×792, slice thickness = 0.24mm, slices per slab = 640, phase encoding direction right to left, slice oversampling = 80%, flip angle = 30deg, TR = 36ms, TA_base_ = 09:08:07h. Supplementary Table S1 summarizes the corresponding acquisition times again and also informs about repeated acquisitions for later averaging.

### Elective postprocessing of image data

For image reconstruction, the standard postprocessing pipeline on the MR system was used, consisting of the basic Fast Fourier Transform (FFT) and the standard intensity normalization filter (vendor name “prescan normalize”). Neither other kinds of filtering nor interpolation algorithms were utilized. The images were exported into the standard DICOM format.

The repeated volume acquisitions of identical resolution were manually averaged up slice by slice directly on the MR system and were then also exported as DICOM images. No kinds of registration procedures were performed.

The presented full-size and zoomed MR images in all figures were generated with the free software ITK-SNAP 3.6.0 ^18^.

### Expert reading

Recommendations for the minimum number of averages (averages_min-SNR_, Supplementary Tab. S1) that should be performed for a given resolution are based on the expertise of seven MRI experienced clinicians and scientists (experience for 12.1 ± 7.8 years, mean ± standard deviation). All of them took part in a small survey by evaluating respective series of images with increasing averages, judging about the displayed SNR and overall image quality.

## Results

Based on representative imaging examples in transverse, sagittal, and coronal orientation, Figs. 1 to 8 as well as Supplementary Figs. S1 to S5 depict the characteristics of the suggested *ex vivo* 3T URI-FLASH approach. As it can be deduced from all the figures, the complete setup of fixed brain, dedicated container, hardware, and MR sequence is stable enough for measuring isotropic ultra-high resolutions up to 160μm in 3D with a pronounced tissue contrast over more than 90h (cf. Supplementary Tab. S1).

**Figure 1:**
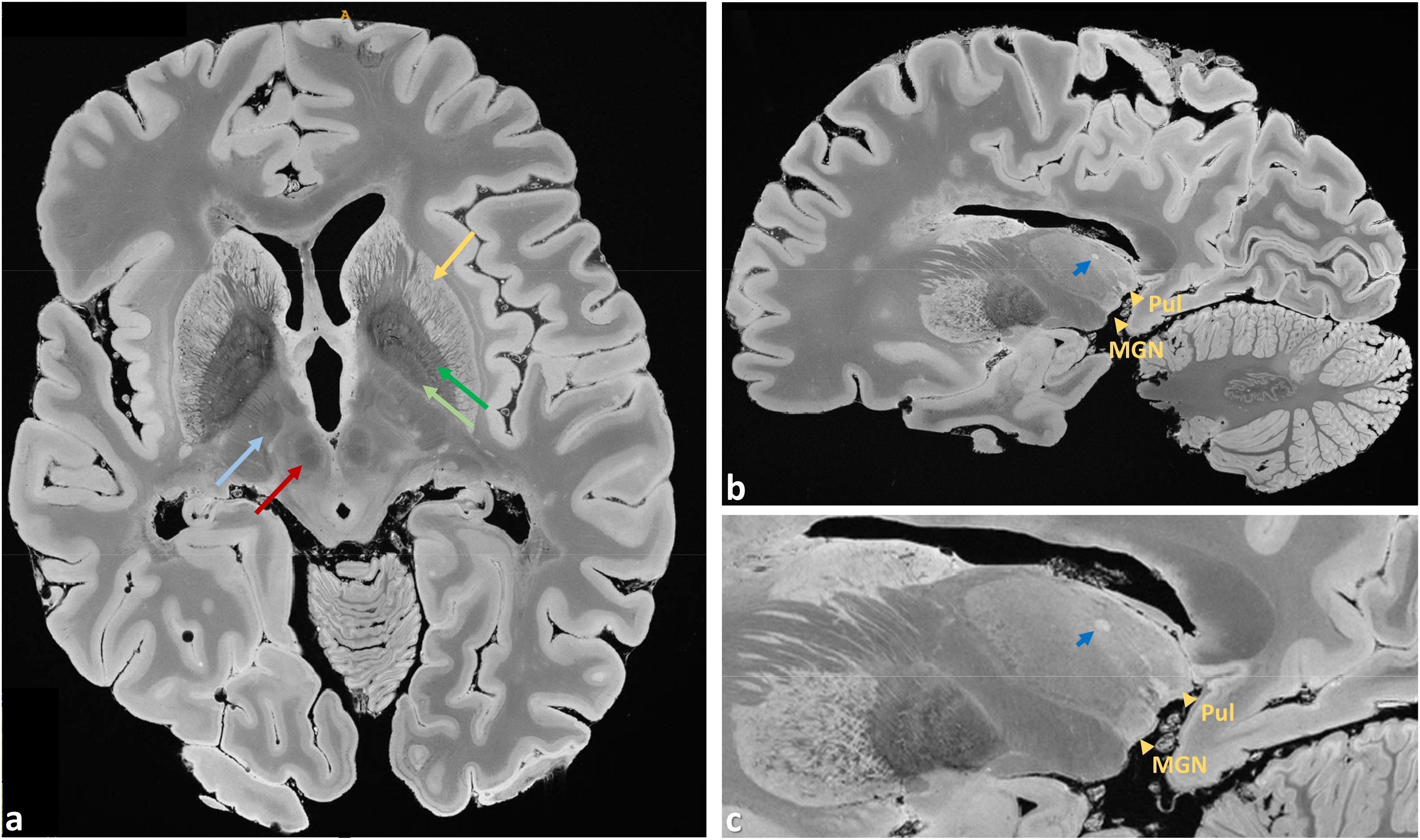
Representative images from the 200μm acquisition demonstrate the richness of anatomical details that can be observed in the URI-FLASH images. They display a combination of resolving capability and soft tissue contrast that can just not be obtained with clinical 3T imaging. **a**: Originally acquired transverse slice depicting, e.g., the putamen (gold arrow), external globus pallidus (dark green arrow), internal globus pallidus (light green arrow), red nucleus (red arrow), and subthalamic nucleus (light blue arrow). **b**: Sagittal reformation and **c**: enlargement of the thalamus and the basal ganglia. Gold arrowheads mark the thalamic nuclei medial geniculate nucleus (*Med Gen*) and pulvinar (*Pul*). A thalamic lesion can be also observed (dark blue arrow).

Generally, a common goal of postmortem MRI is the maximization of the acquired spatial resolution. Using the example of the dentate nucleus, Fig. 2 points out that some structures in the human brain are so hyperfine that even an ultra-high-resolution gain from 240μm towards 160μm is still valuable. Supplementary Figure S1 also shows a comparison for the line of Gennari. The ultra-high-resolution acquisitions only make sense, however, if the targeted resolution can be realized with “sufficient SNR” within the allotted total acquisition time. Supplementary Table S1 contrasts the minimum total acquisition times (i.e., averages) needed for a desired spatial resolution to obtain such a “sufficient SNR”, as perceived in the eyes of seven clinicians and scientists having long experience with MRI. Supplementary Table S1 also presents the maximum acquisition times per resolution that could be realized within the frame of this work. As a further noteworthy side information, the relative SNRs of the protocols are given.

**Figure 2:**
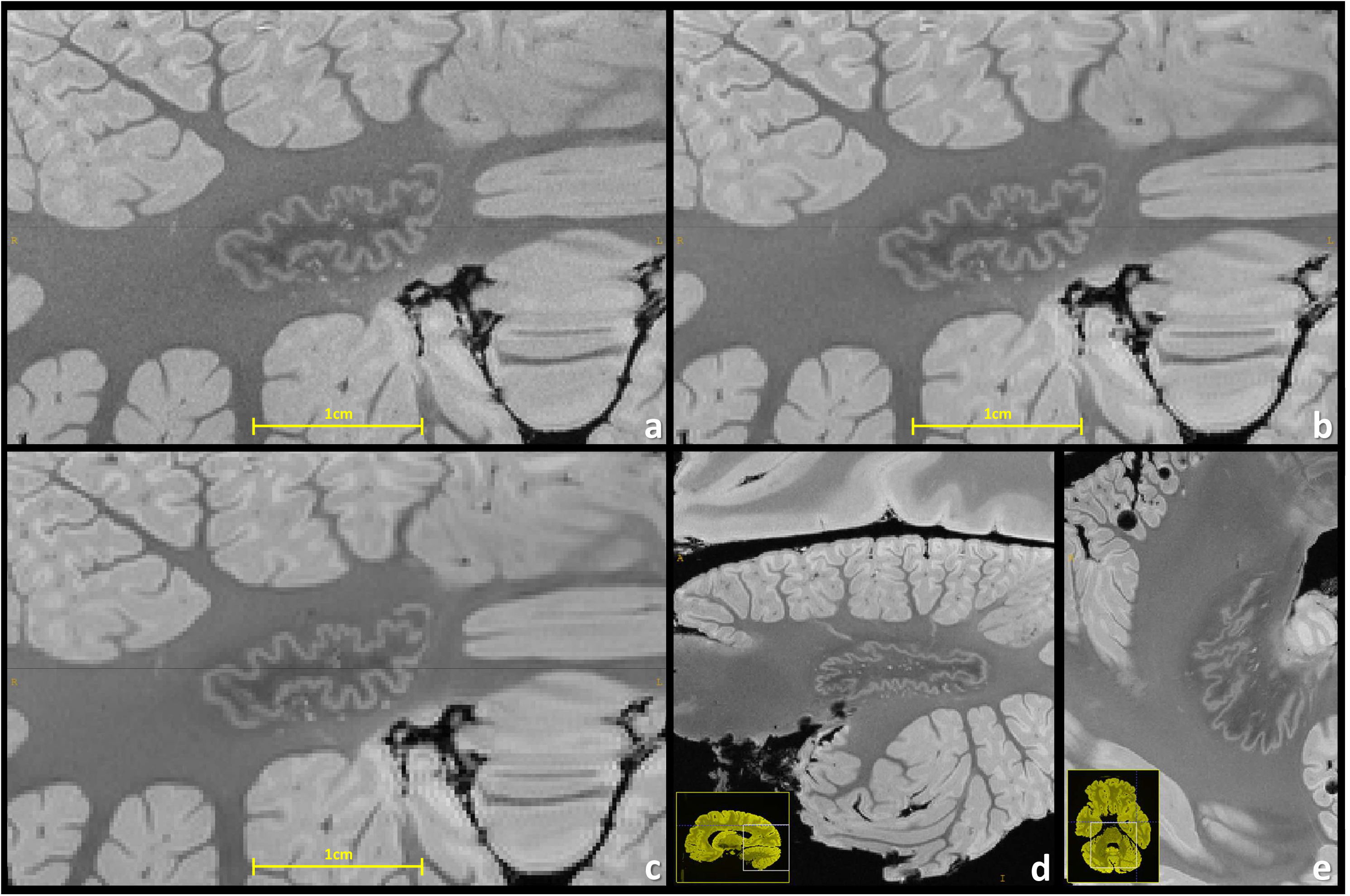
Comparison of resolving power visualized at the cerebellar dentate nucleus with 160μm (**a**), 200μm (**b**), and 240μm (**c**) resolution. Particularly for the 240μm resolution the structures get pixelated, demonstrating that there is still gain in increasing even such ultra-high resolutions for brain MRI. A sagittal reformation (**d**) and a transverse slice (**e**) of the dentate nucleus are also depicted (200μm resolution).

The individual results of the “minimum SNR survey” are presented in Supplementary Tab. S2. Supplementary Figure S2 provides examples of images at different spatial resolution: acquisitions with the four different base protocols of 240μm, 200μm, 180μm, and 160μm are contrasted with their corresponding averaged acquisitions with both “minimum SNR” and “maximum performed”.

Figures 3 to 8 display a representative compilation of different small-sized multiple sclerosis lesions (purely intracortical and leukocortical lesions, Figs. 3 to 5; lesions with central vein, Figs. 3 and 5; lesions with hypointense rims, Fig. 4; subpial lesions, Fig. 6; hypointense juxtacortical rims, Fig. 6; thalamic and subcortical WM lesions, Fig. 7; lesions in both the dentate nucleus and the olivary nucleus in the pons, Fig. 8). These exemplary lesions clearly show that a combination of a good SNR, very-high spatial resolution and strong tissue contrast is essential for delineating these fine structures. We were able to identify lesions with peculiar hypointense patterns (open and closed rims within the cortex and in the juxtacortical areas) as well as lesions in the grey matter nuclei of the brainstem (i.e. olivary nucleus).

**Figure 3:**
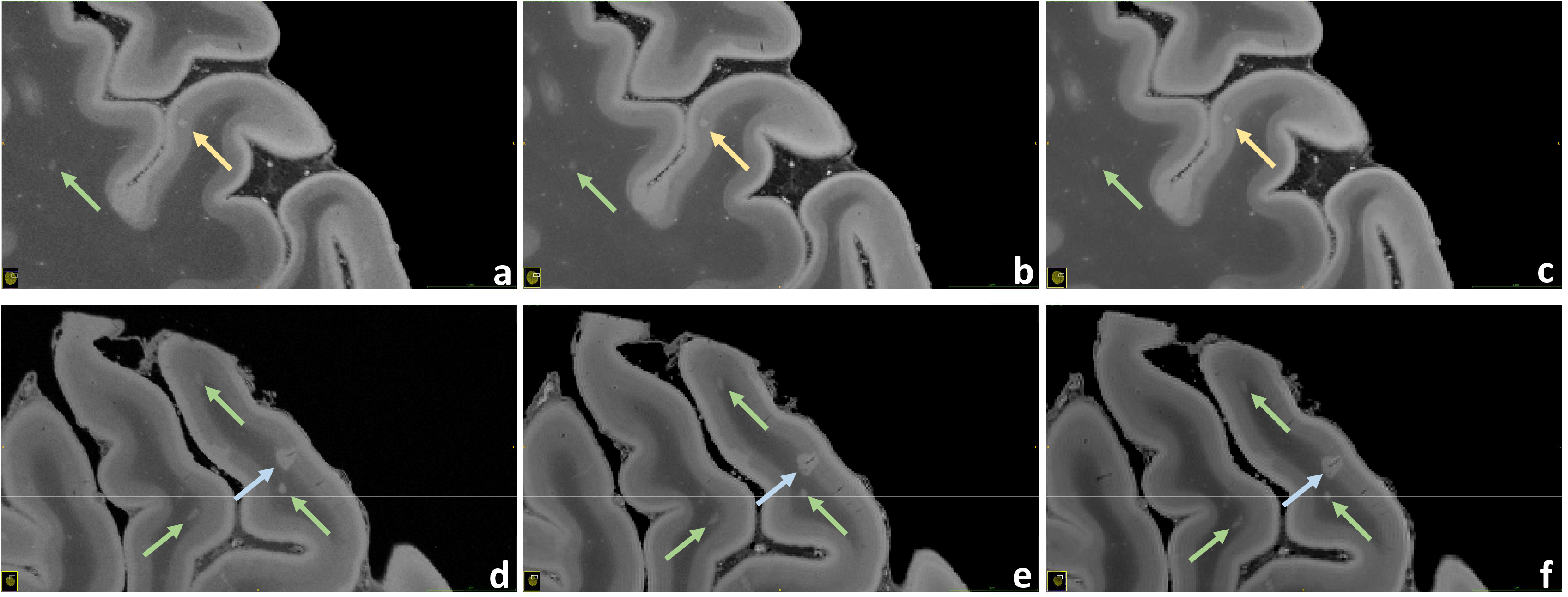
Juxtaposition of intracortical, leukocortical, and WM lesions. *Upper row*: A pure intracortical lesion acquired with 160μm (**a**), 200μm (**b**) and 240μm (**c**) resolution is shown (gold arrows). This very small round hyperintense area is placed in the middle of the cortex, without touching the WM or the subpial surface. *Lower row*: A leukocortical lesion acquired with 160μm (**d**), 200μm (**e**) and 240μm (**f**) resolution is depicted (blue arrows). In this case the hyperintense area is located across the border between WM and GM. A thin linear hypointensity, representing the “central vein”, is well visible. These scans also enable the visualization of very small lesions in the WM, probably representing nascent WM pathology (green arrows).

**Figure 4:**
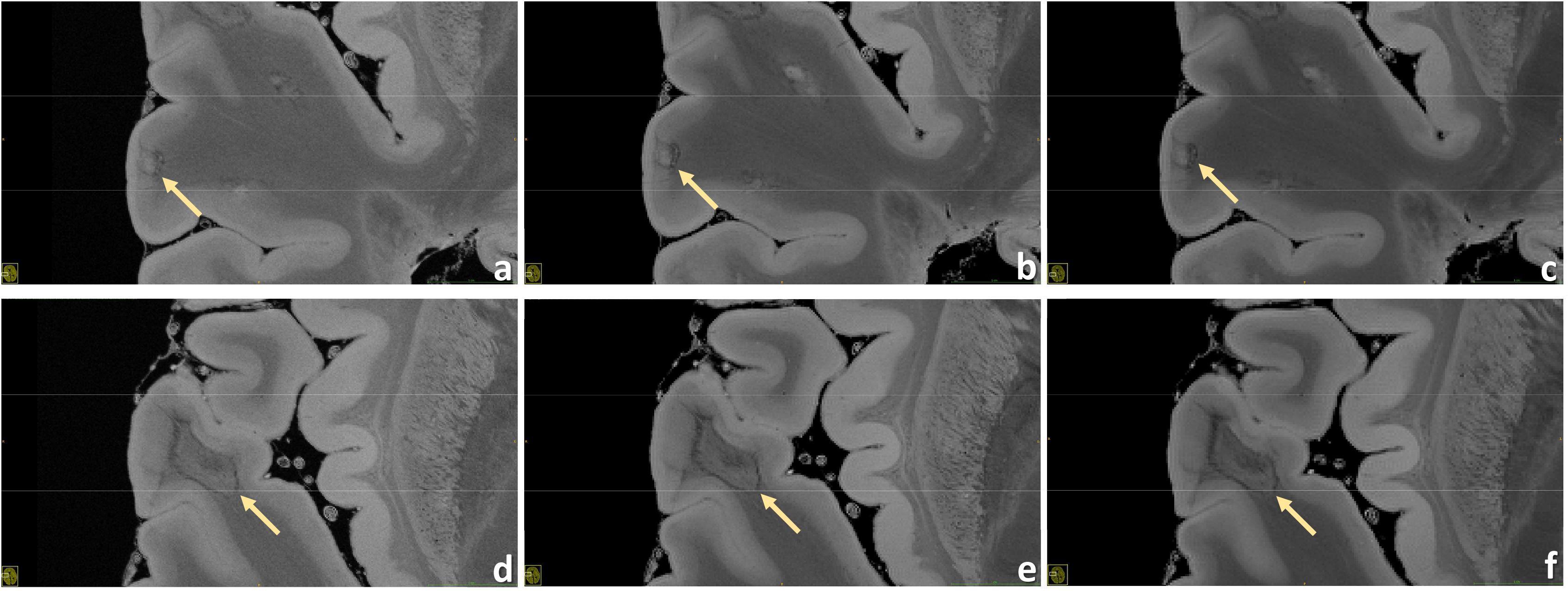
Two different leukocortical lesions are contrasted. *Upper row*: A leukocortical lesion with hypointense “rim” at 160μm (**a**), 200μm (**b**) and 240μm (**c**) resolution is displayed. This inhomogeneous area located across the border between WM and GM presents a central hyperintensity surrounded by a hypointense frame. *Lower row*: A leukocortical lesion with hypointense “rim” limited at the WM part at 160μm (**d**), 200μm (**e**) and 240μm (**f**) resolution is demonstrated. In this case the hyperintense WM part of the lesion is surrounded by an hypointense frame; in the nearby cortex a light focal hyperintensity can be noticed.

**Figure 5:**
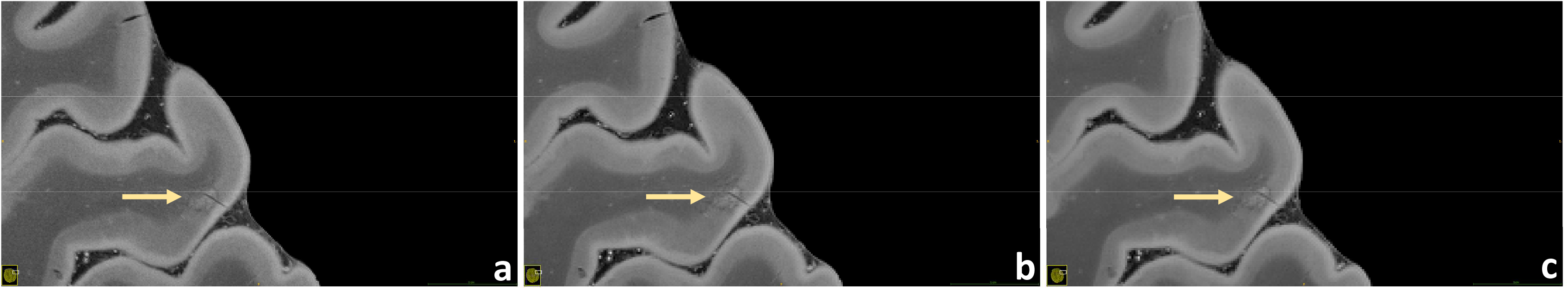
A leukocortical lesion with “central vein” acquired at 160μm (**a**), 200μm (**b**) and 240μm (**c**) resolution is presented. This inhomogeneous area is almost completely contained in the cortex and presents hypointense borders. The “central vein”, a threadlike hypointensity in the middle of the lesion, can be observed easily.

**Figure 6:**
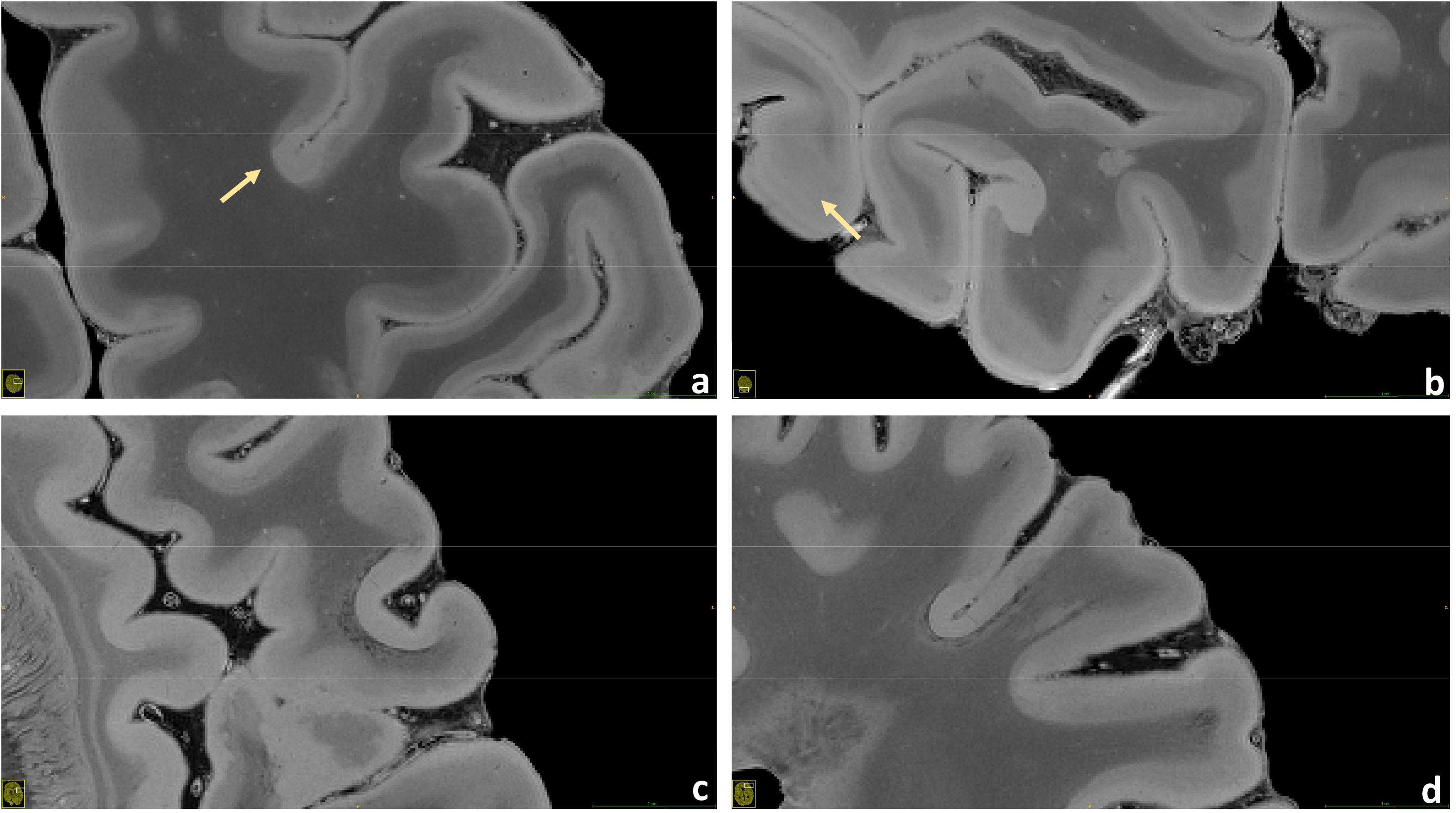
A small potpourri of four putative subpial lesions measured at 160μm (**a**, **b**) and 200μm (**c**, **d**) resolution is illustrated. In this sequence the cortical structure mostly appears bilayered, probably expressing the different myelin and iron content between the external and the internal layers of the cortex. However, focal losses of this structural organization can be observed well in the pictures: areas, which do not follow the irroration territory of cortical veins and are especially found in proximity of the sulci, present hyperintense throughout the whole cortex with loss of the otherwise well visible bilayered structure. These pathoanatomical changes most probably represent subpial lesions. Some of these lesions show a hypointense juxtacortical rim (**c**, **d**).

**Figure 7:**
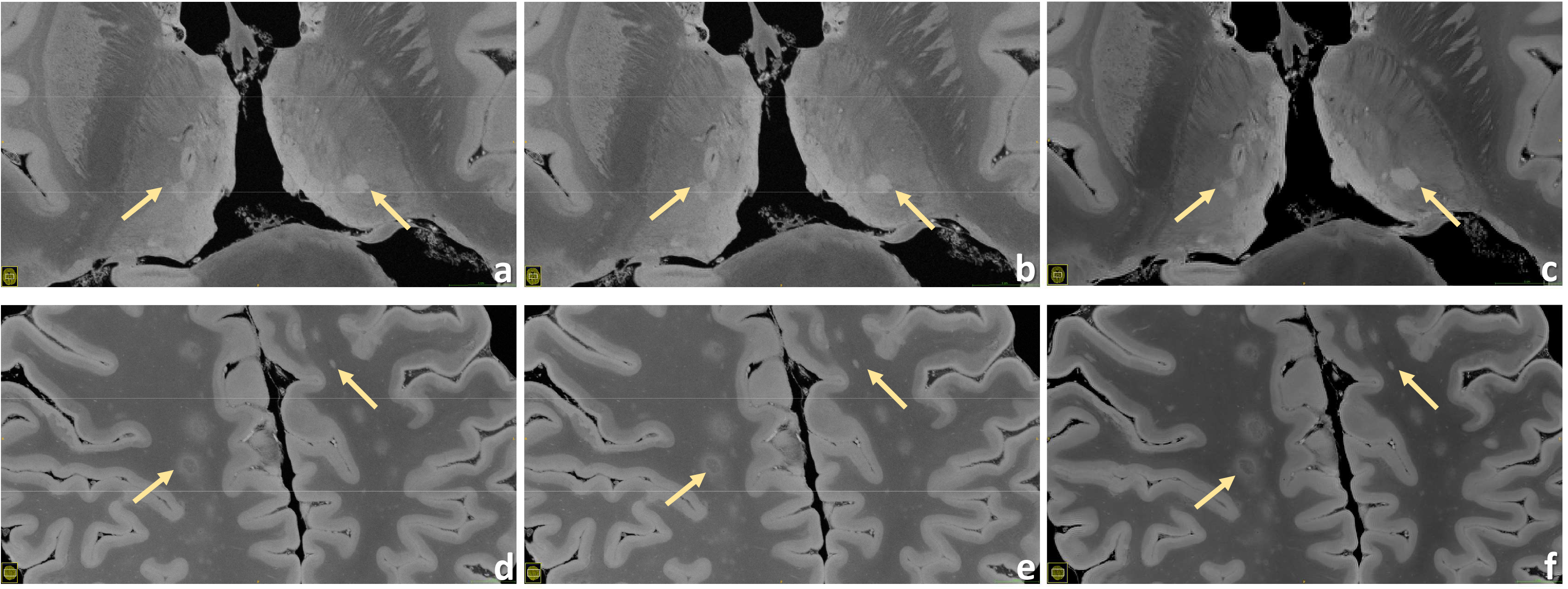
Examples of thalamic lesions (*upper row*) and subcortical WM lesions (*lower row*) are demonstrated for the 160μm (**a**, **d**), 200μm (**b**, **e**) and 240μm (**c**, **f**) resolution.

**Figure 8:**
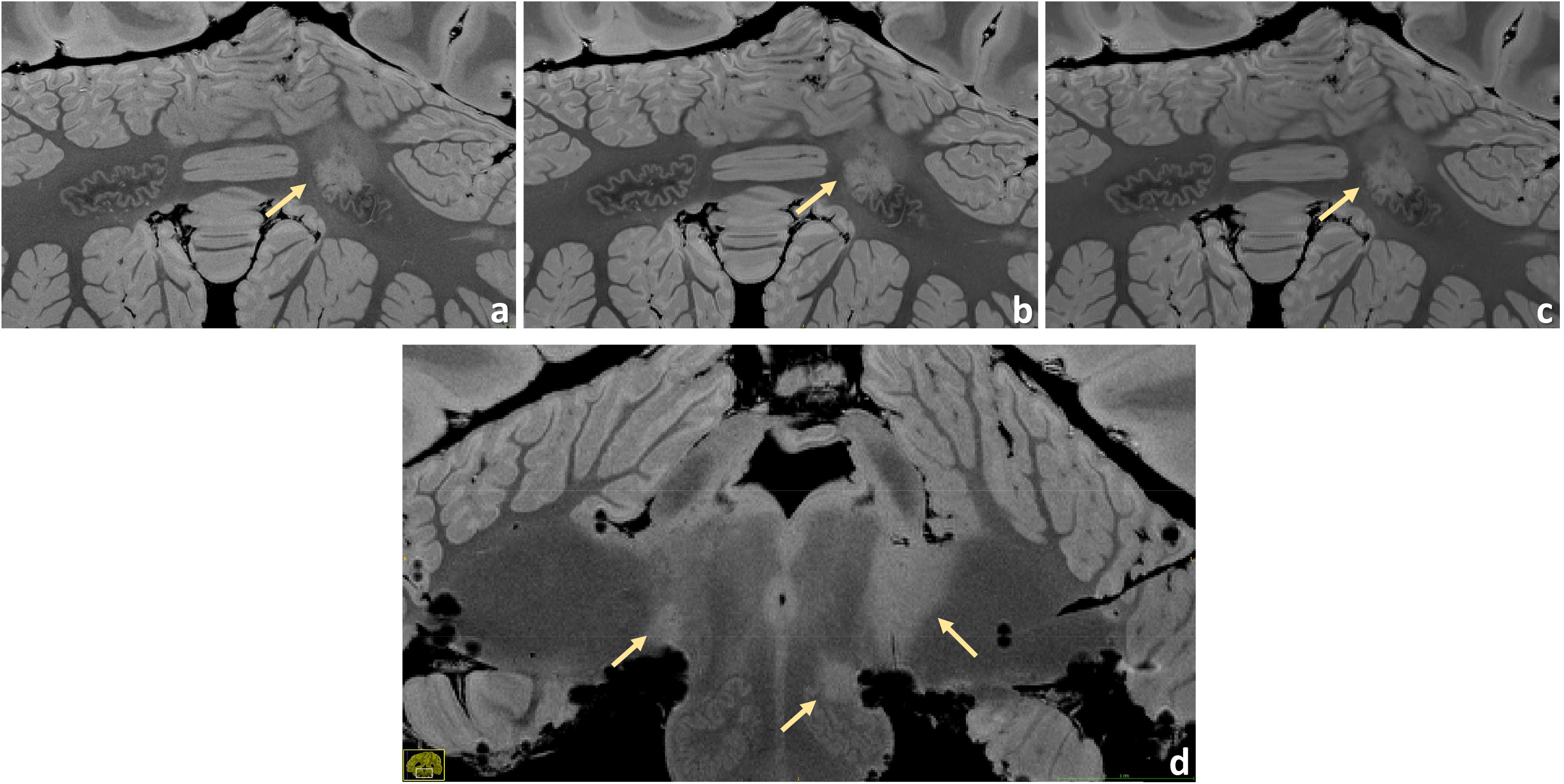
Examples of infratentorial lesions. A dentate nucleus lesion is contrasted at the 160μm (**a**), 200μm (**b**) and 240μm (**c**) resolution. The anatomical features of this nucleus are depicted in great detail, especially at larger magnification. In this case, the left dentate nucleus is lesioned in its rostral part. A very detailed anatomical representation of brainstem and cerebellum is shown at 160 μm resolution (**d**). A small lesion of the left inferior olivary nucleus is well visible and clearly defined. Moreover, typical bilateral MS lesions at the cerebellopontine angle can be observed here, more extended on the left side.

Altogether for Figs. 2-5, 7 and 8 as well as for Supplementary Fig. S1, it should be noted how very well equivalent slices of similar geometry could be found in the datasets of different ultra-high resolutions, even though the brain container was completely removed from the MR system in between: the acquisitions were independent with at least one week apart.

## Discussion

### Technical and methodological aspects

We developed an *ex vivo* whole-brain MR imaging approach that facilitates isotropic 3D ultra-high-resolution imaging up to 160μm with a strong soft tissue contrast on a common clinical 3T MR system. By utilizing only standard hardware components, the rationale was to test the viable boundaries for *ex vivo* URI at 3T and, consequently, establishing an MRI concept that can be well reproduced on other MR systems. Thus, our approach was substantially different from that of a recently published work^4^, where a spatial resolution of 100μm was achieved in a brain specimen; the latter employed a similar acquisition sequence but made use of 7T field strength, of a custom-made phased-array coil as well as of self-made computational tools that allow bypassing the raw data storage capacity of the MR system, did offline MRI reconstruction and enable post-processing of a few terabytes of MR raw data ^4^.

Four base protocols of 160μm, 180μm, 200μm, and 240μm isotropic resolution were established. Acquisition times between approximately 7 hours up to 90 hours were investigated. Even the ‘low’ 240μm base protocol offers great potential for 9h acquisition time; this protocol can be easily added to other postmortem examinations. As our data show, the full capabilities of a 160μm acquisition are revealed for our *ex vivo* approach covering at least ca. 42 hours – the recommended minimum acquisition time based on Supporting Tab. S1 –, which can be usually organized well in the light of routine measurements at site by utilizing weekends.

Generally, the presented *ex vivo* URI-FLASH approach is based on four major reasonings: (1) we experienced that RF spoiled gradient-echo sequences, which are also harnessed for quantitative susceptibility mapping ^19^, facilitate acquiring high-quality MR images with remarkable soft-tissue contrast under *ex vivo* conditions; (2) the T1 relaxation times are considerably shortened in fixed *ex vivo* tissue ^20^, which leads to a much faster recovery of longitudinal magnetization. RF spoiled gradient-echo sequences with short repetition time TR, demonstrating steady state magnetization response ^21,22^, benefit a lot from this effect: they can be tuned to a much higher signal efficiency compared to *in vivo* conditions; (3) an exceptionally low receiver bandwidth of 50Hz/Pixel was used to enhance considerably the overall SNR and optimize the time the MR systems spends on signal acquisition for the given echo time TE and repetition time TR. Additionally, it enabled comparatively low gradient strengths and moments for high spatial resolutions. (4) The full value for basic and clinical neuroscience is obtained for isotropic 3D resolutions alone: only then an unbiased view is possible, effects like partial volume are invariant of the direction, and multi-planar reformations are feasible in arbitrary directions without loss of information.

A recent publication presents a SPACE sequence based approach with a 400μm isotropic resolution acquired on a 3T system ^5^. The work already suggests the principal value of acquiring very-high isotropic spatial resolutions at 3T. Our URI-FLASH based work achieves an unprecedented 160μm spatial resolution for whole brain imaging at 3T. Nevertheless, acquiring a (400/160)^3^ = 15.6 times smaller voxel volume compared to Ref. ^5^ leads to a 15.6 times lower SNR accordingly. Just compensating this drastic SNR loss by signal averaging would lead to a 15.6^2^ = 244 times longer acquisition time. Although this rough calculation is just volume-based and neglects spatial encoding, it already shows that the presented URI-FLASH approach is much more efficient in terms of SNR. The main reason for this fact are rationale (2) and (3). On the contrary, for the SPACE ^23^ sequence, being a specialized 3D variant of the turbo spin echo sequence with low refocusing flip angles ^24^, the considerably shortened T1 and T2 relaxation times under *ex vivo* conditions are of clear disadvantage regarding acquisition efficiency and also regarding image quality in terms of image blurring ^5,25^. Moreover, receiver bandwidths as low as 50Hz/Pixel would have a further detrimental effect on the SPACE sequence’s image quality.

As already noted above, increasing the isotropic 3D spatial resolution in MRI comes at the expense of considerable SNR reduction that can be only compensated for by signal averaging in our experiments. Considering the general ‘square root law’ between SNR and acquisition time, signal averaging is an inefficient procedure to increase SNR for higher numbers of averages. Consequently, higher resolutions lead to a dramatic prolongation of acquisition times. Thus, on one hand, it may be worthwhile to limit the number of averages to the absolute minimum. On the other hand, even a moderate noise impact may hamper the resolving of very fine structures, as it can be found in the basal ganglia or in the cerebellum. To investigate this effect for the four different spatial resolutions, a small survey was organized: seven clinicians and scientists with experience in brain MRI evaluated respective series of images with increasing averages, judging about the displayed SNR and overall image quality. As a result, resolution-specific recommendations for a minimum of averages (i.e., minimum total acquisition time) perceived to be essential for “sufficient SNR” were defined (Supplementary Tab. S1, individual decisions in Supplementary Tab. S2). A visual comparison of the resulting decisions is presented in Supplementary Fig. S2. With the aid of Supplementary Tab. S1 and Supplementary Fig. S2 the reader can orientate on which resolution protocol to choose for a given, available total acquisition time. Moreover, Figs. 2-5, 7 and 8 as well as Supplementary Fig. S1 combine images of different spatial resolutions such that the reader can get a better notion of possible differences and what image quality can be expected for a given acquisition time, but also presenting a variety of locations and lesions at the same time.

Manufacturer-supplied MR sequences limit which MR parameters can be influenced and particularly their range of values. *Ex vivo* MRI often requires parameter combinations and values, however, which are uncommon in clinical *in vivo* MRI. Consequently, we programmed a basic RF spoiled gradient echo (FLASH) sequence that allows broad parameter ranges and bypasses part of the internal software like view-ordering schemes. As a result, we could avoid non-desirable limitations for parameters such as base matrix size and maximal product of phase encoding steps in 3D. Additionally, it was made sure that all gradient durations were stretched to the full possible limit, thereby reducing the gradient amplitude (‘natural’ exclusions: readout and slab selection gradient). Thus, our URI-FLASH approach was eventually limited by the maximal image reconstruction size supported by the MR system. To avoid any conflict with the limited raw data storage space on the MR system, single acquisitions were performed first and averaged up manually at a later timepoint (cf. Supplementary Fig. S2).

The targeted resolutions require acquisition times of hours to days on a clinical MR system, where the whole gradient and RF hardware components may be permanently operated at maximum performance: this situation represents an extreme strain on the hardware components, which can even lead to their damage or failure. Moreover, factors like maximum gradient strength, maximum gradient power amplifier duty cycle, and maximum RF power amplifier duty cycle put restrictions to the maximal resolution and acquisition efficiency. A substantial benefit of the used URI-FLASH approach is that its strain on the hardware is low despite the requirements: the exceptionally low bandwidth leads to relatively weak gradients and gradient moments on the readout axis. Furthermore, with TR/2 ≅ TE = 18ms, enough time is left for the phase and 3D encoding gradients in the sequence. Altogether, this facilitates the ultra-high resolutions we could achieve and holds mechanical vibrations in the MR system at bay. Naturally, strong mechanical vibrations affect URI adversely and can even stimulate increased helium boil-off if the mechanical vibrations hit certain eigenfrequencies of the MR system.

Similarly, the strain on the RF hardware is also in the normal range for URI-FLASH, since only one RF excitation pulse of low flip angle every TR ≈ 37ms is applied. To maximize the tissue SNR, the excitation flip angle was chosen close to the expected Ernst angle ^26^ and, hence, depends on the TR (cf. Methods section).

In line with our general rationale of establishing a reproducible *ex vivo* URI approach on a common 3T MR system, we employed the standard 20-channel phased-array head and neck coil supplied by the manufacturer. On the one hand, the spaciousness of this coil helps to accommodate the ad-hoc brain container. Moreover, it provides a quite homogeneous B_1_ reception field, which - in combination with the quite homogeneous B_1_^+^ excitation field of the 3T MR system - leads to MR images of overall homogeneous tissue intensity and contrast. A comparison of Figs. 1 to 4 with images of Ref. ^4^ substantiate this observation. A further advantage is that fewer coil elements necessitate less reconstruction memory on the MR system; thus, since we utilized all the available MR scanner host resources, the maximal resolution that could be reconstructed depended on the number of coil elements. On the other hand, a drawback of the 20-channel head and neck coil is the notably lower SNR achieved compared to an alternative vendor coil with more coil elements or a custom-made coil.

### Clinical aspects

Figures 1 to 8 and Supplementary Figs. S1 to S5 all underline the potential of 3T based *ex vivo* examinations in clinical neuroscience.

First, URI-FLASH allows to distinguish different layers of the cerebral cortex, namely, the external part of the cortex appears more hyperintense, most probably representing the subpial molecular layer and the external part of the granular layer (Figs. 3 to 6). Furthermore, the line of Gennari, in the calcarine fissure, is hypointense (Supplementary Fig. S1). Second, because of its tissue-sensitive contrast and the sensitivity to susceptibility effects, URI-FLASH also permits the identification of some intrathalamic nuclei like the pulvinar nuclei and the medial geniculate nucleus (Fig. 1 b,c) as well as the medio dorsal nucleus, anterior nucleus and the *lamina intermedia* (Supplementary Fig. S3).

Since URI-FLASH enables a fine depiction of the intracortical structure, it permits an easier identification of subpial lesions (Fig. 6), which are the most specific and frequent cortical lesion subtype in MS patients^27,28^ – but which are barely recognizable in *in vivo* 3T MRI. Identifying cortical demyelinated lesions with MRI is challenging due to the small change in myelin-related signal that is observed in those lesions compared to WM lesions. Besides, cortical lesions are often less inflammatory than WM lesions ^29^, leading to limited alteration of signal intensity in T1 and T2 weighted MRI ^30,31^. URI-FLASH at 3T shows therefore not only the advantage of a very high micrometric spatial resolution but also to the subtle demyelinating process occurring in cortical lesions in MS patients, as previously shown at 7T MRI ^32^.

Besides, the contrast provided by URI-FLASH – which combines an exquisite sensitivity to the presence of paramagnetic substances such as iron – allows to identify multiple characteristics of both cortical and white matter MS lesions, such as the presence of complete iron rims surrounding leukocortical lesions (Fig. 4 a,b,c) or just their white matter part (Fig. 4 d,e,f). Lesions harboring paramagnetic rims, which are reflecting the accumulation of microglia cells and ongoing smoldering inflammatory activity ^33,34^, are particularly important as they have been linked to disease progression and high disability in MS patients ^33,35^. Hence, URI-FLASH may provide a new window to understand the role of those lesions in the cortical layers and in the juxtacortical white matter.

Likewise, URI-FLASH provides evidence of gradients of tissue pathology, both in WM and in GM, which are clearly paralleling the patterns described in neuropathological studies ^27,36,37^. The micrometric resolution of URI-FLASH images also provides a clear advantage for the study of nascent sub-millimetric pathology in WM (Fig. 3), small punctiform lesions in cortical GM (Fig. 3) and lesions affecting the cerebellar nuclei and the convoluted GM/WM cortical layers of the cerebellar cortex (Fig. 8) and the deep GM nuclei (Fig. 1 and Supplementary Fig. S3).

To conclude, URI-FLASH provides a microscopic insight of the whole-human brain at 3T, which is achieved through a micrometric resolution, a tissue-specific contrast and a superb sensitivity to susceptibility effects. URI-FLASH is therefore an excellent approach to investigate the microscopic characteristics of the human brain as well as its changes in a complex pathology like multiple sclerosis.

## Acknowledgements

This work was funded by the Swiss National Science Funds PP00P3_176984 and supported by the German Ministry of education (BMBF; KKNMS German competence network for multiple sclerosis). Govind Nair is supported by the Intramural Research Program at the NINDS.

## Author contributions

MW: Conceptualization, Methodology, Software, Validation, Investigation, Data Curation, Writing - Original Draft, Writing - Review & Editing, Visualization, Project administration

PD: Validation, Investigation, Data Curation, Instrumentation, Writing - Review & Editing

RG: Methodology, Validation, Data Curation, Writing - Original Draft, Writing - Review & Editing, Visualization

EB: Validation, Investigation, Writing - Review & Editing

GN: Methodology, Software, Writing - Review & Editing;

LK: Validation, Writing - Review & Editing

WB: Resources, Validation, Writing - Review & Editing

CS: Resources, Validation, Writing - Review & Editing

CG: Conceptualization, Validation, Investigation, Resources, Data Curation, Writing - Original Draft, Writing - Review & Editing, Visualization, Supervision, Project administration, Funding acquisition

## Additional Information

## List of abbreviations (alphabetical)

URI: ultra-high-resolution imaging
FLASH: fast low-angle shot
GM: gray matter
WM: white matter

