## Supplemental Tables for "Imaging Multiple Sclerosis Pathology at 160μm Isotropic Resolution by Human Whole-Brain *Ex Vivo* Magnetic Resonance Imaging at 3T"

#### Supporting Table S1: Overview of Acquisition Times for the Investigated Protocols

| **Isotropic 3D resolution:** | **240µm** | **200µm** | **180µm** | **160µm** |
|---|---|---|---|---|
| **$SNR_{relative}$ / a.u.** | 1.00 | 0.54 | 0.36 | 0.25 |
| **$TA_{base}$ / h** | 09:08:07 | 08:12:33 | 07:04:49 | 06:57:20 |
| **$averages_{min-SNR}$ #** | 3 | 4 | 5 | 6 |
| **$TA_{min-SNR}$ / h** | 27:24:21 | 32:50:12 | 35:24:05 | 41:44:00 |
| **$averages_{max-performed}$ #** | 9 | 11 | 8 | 11 |
| **$TA_{max-performed}$ / h** | 82:13:03 | 90:18:03 | 56:38:32 | 76:30:40 |

$SNR_{relative}$: Relative SNR of the different acquisition protocols.

$TA_{base}$: Acquisition time of the base protocol (single acquisition without averaging)

$averages_{min-SNR}$: Minimum number of averages recommended for "sufficient SNR".

$TA_{min-SNR}$: Resulting acquisition time for the recommended "minimum SNR protocol"

$averages_{max-performed}$: Maximum number of averages that could be performed within the frame of the investigations.

$TA_{max-performed}$: Resulting acquisition time for maximum number of averages performed.

[#] Manual averaging of the acquired magnitude images from repeated base protocol measurements.

#### Supporting Table S2: Overview of Expert Decisions for the Visual SNR Evaluation

| **Recommendations:** | **240µm** | **200µm** | **180µm** | **160µm** | **Expertise** |
|---|---|---|---|---|---|
| **$\text{averages}_{\text{min-SNR}}$ (E1)** # | 1 | 3 | 3 | 5 | Neurologist |
| **$\text{averages}_{\text{min-SNR}}$ (E2)** # | 6 | 4 | 5 | 6 | Neuroradiologist |
| **$\text{averages}_{\text{min-SNR}}$ (E3)** # | 1 | 2 | 2 | 2 | Neurologist |
| **$\text{averages}_{\text{min-SNR}}$ (E4)** # | 4 | 5 | 7 | 6 | Image Processing Expert |
| **$\text{averages}_{\text{min-SNR}}$ (E5)** # | 3 | 6 | 6 | 9 | MRI specialist (Biologist) |
| **$\text{averages}_{\text{min-SNR}}$ (E6)** # | 1 | 1 | 3 | 4 | Neurologist |
| **$\text{averages}_{\text{min-SNR}}$ (E7)** # | 3 | 5 | 6 | 8 | Physicist |
| **arithmetic MEAN** | **2.7** | **3.7** | **4.6** | **5.7** | --- |
| **MEDIAN** | **3** | **4** | **5** | **6** | --- |

$\text{averages}_{\text{min-SNR}}$: Minimum number of averages recommended for "sufficient SNR", here, listed by expert-rater who intentionally have different expertise, i.e., different clinical and scientific backgrounds.

### Manual averaging of the acquired magnitude images from repeated base protocol measurements
