## Supplemental Figures for "Imaging Multiple Sclerosis Pathology at 160μm Isotropic Resolution by Human Whole-Brain *Ex Vivo* Magnetic Resonance Imaging at 3T"

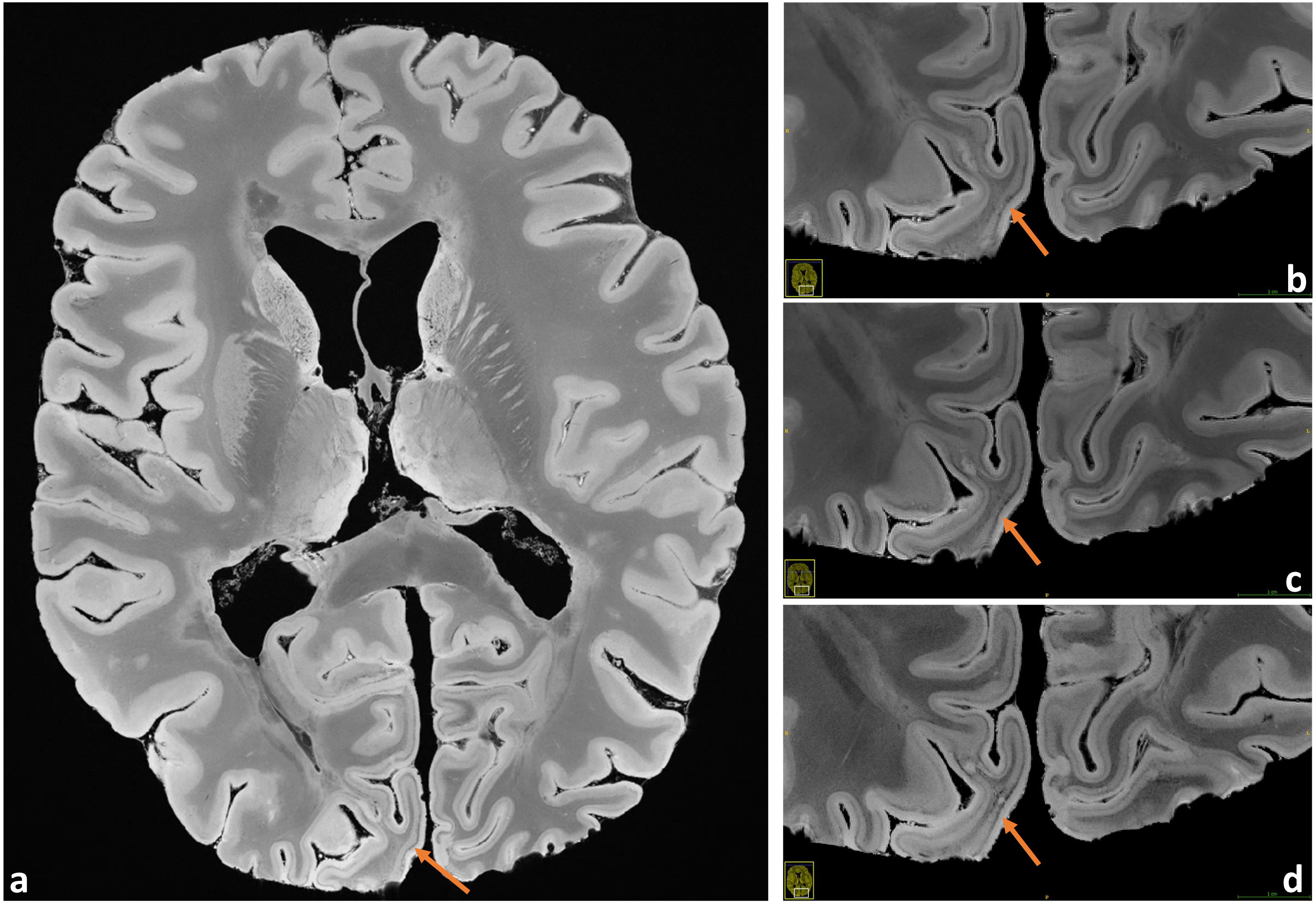


**Supporting Figure S1:** Very good observation of the line of Gennari in the occipital lobe (orange arrows).
**a**: Transverse overview slice, also underlining the pronounced depiction of soft tissue structures; magnifications for the spatial resolutions 240µm (**b**), 200µm (**c**), and 160µm (**d**) are contrasted.

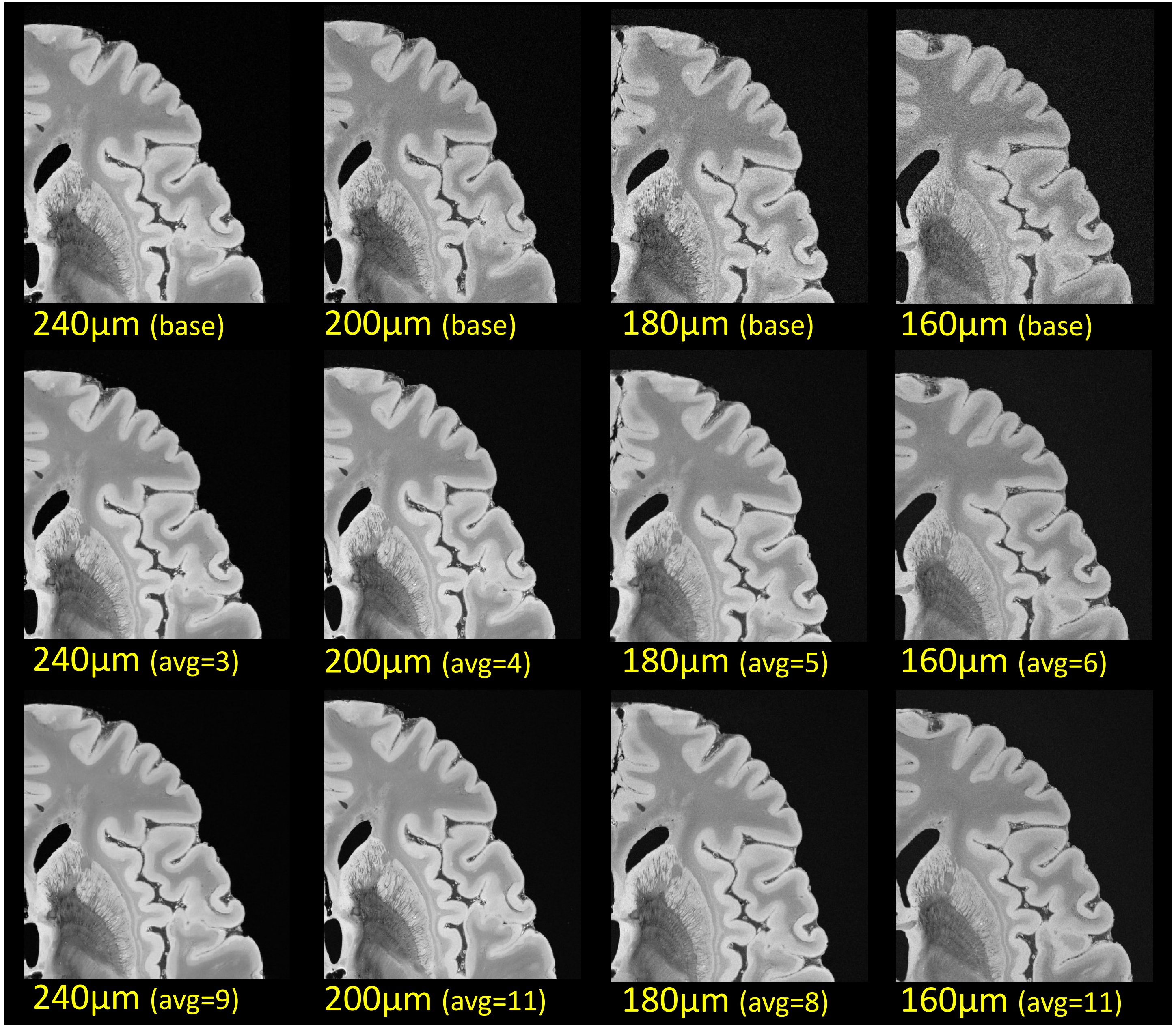


**Supporting Figure S2:** Contrasting juxtaposition of the four acquired resolutions and the effect of averaging. Generally, only quarter images are presented to improve the recognition of SNR changes; the window-levelling is intentionally 'bright'.

*Upper row*: Display of the single acquisitions as specified (base protocol, i.e., no averaging).

*Middle row:* Averaged acquisitions with a "minimum number of averages" to obtain "sufficient SNR for investigation and diagnosis" at the respective resolution, based on a survey under experienced clinicians and scientists, see Suppl. Tabs. S1 and S2.

*Lower row:* Corresponding images with the maximal number of averages that could be performed within the frame of this work.

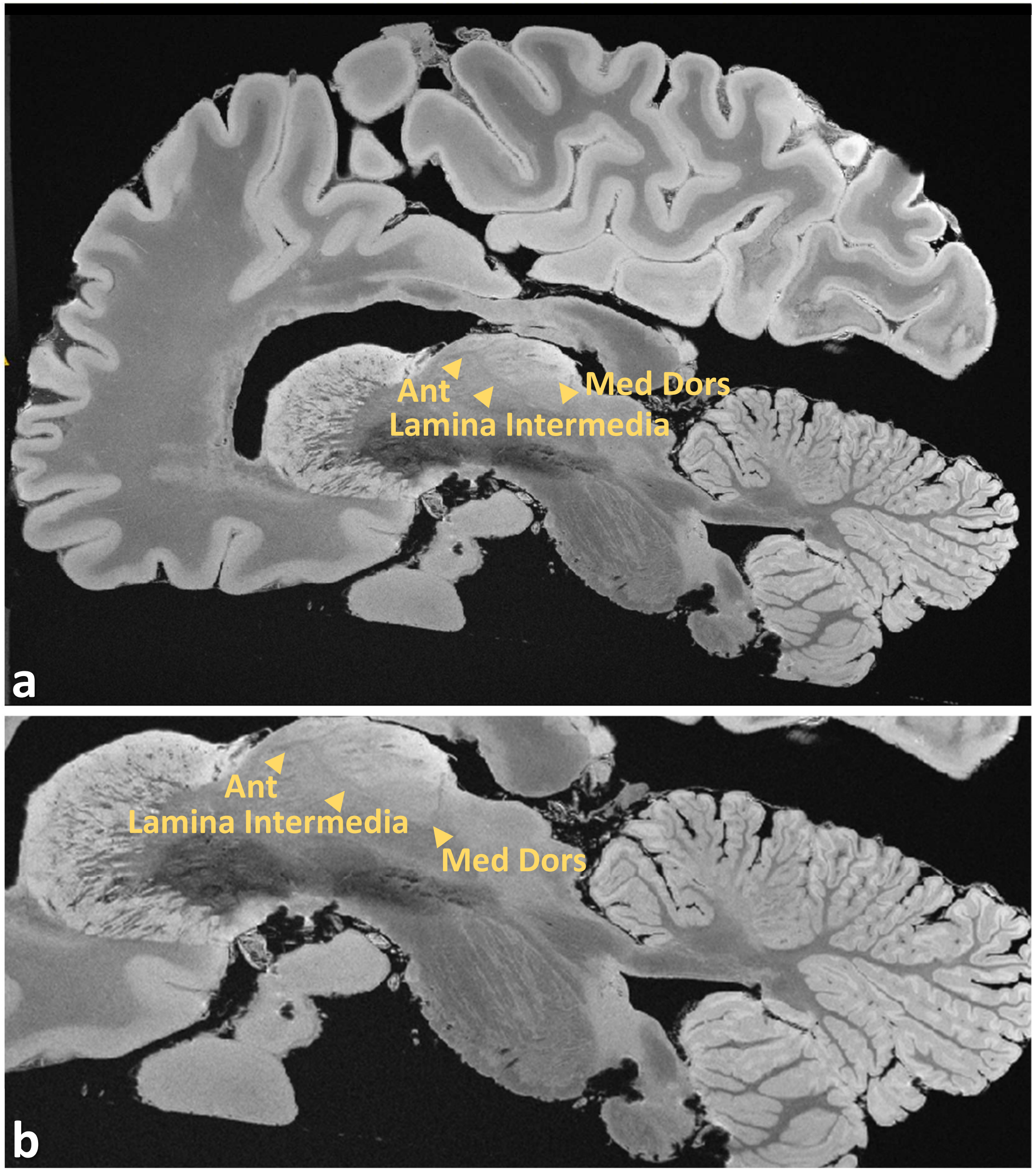


**Supporting Figure S3:** Not least due to the full isotropic 3D resolutions, the sagittal reformations of the MRI datasets also reveal fascinating details. A 160µm cross-section (**a**) with an enlargement of the basal ganglia, the thalamus with its nuclei, the brainstem and the cerebellum (**b**) are shown. Gold arrowheads mark the thalamic nuclei as follows: medio dorsal (*Med Dors*), anterior (*Ant*), *lamina intermedia*, and medial geniculate nucleus.
The two subfigures also display in unprecedented detail the cerebellum's *arbor vitae* at 3T field strength.

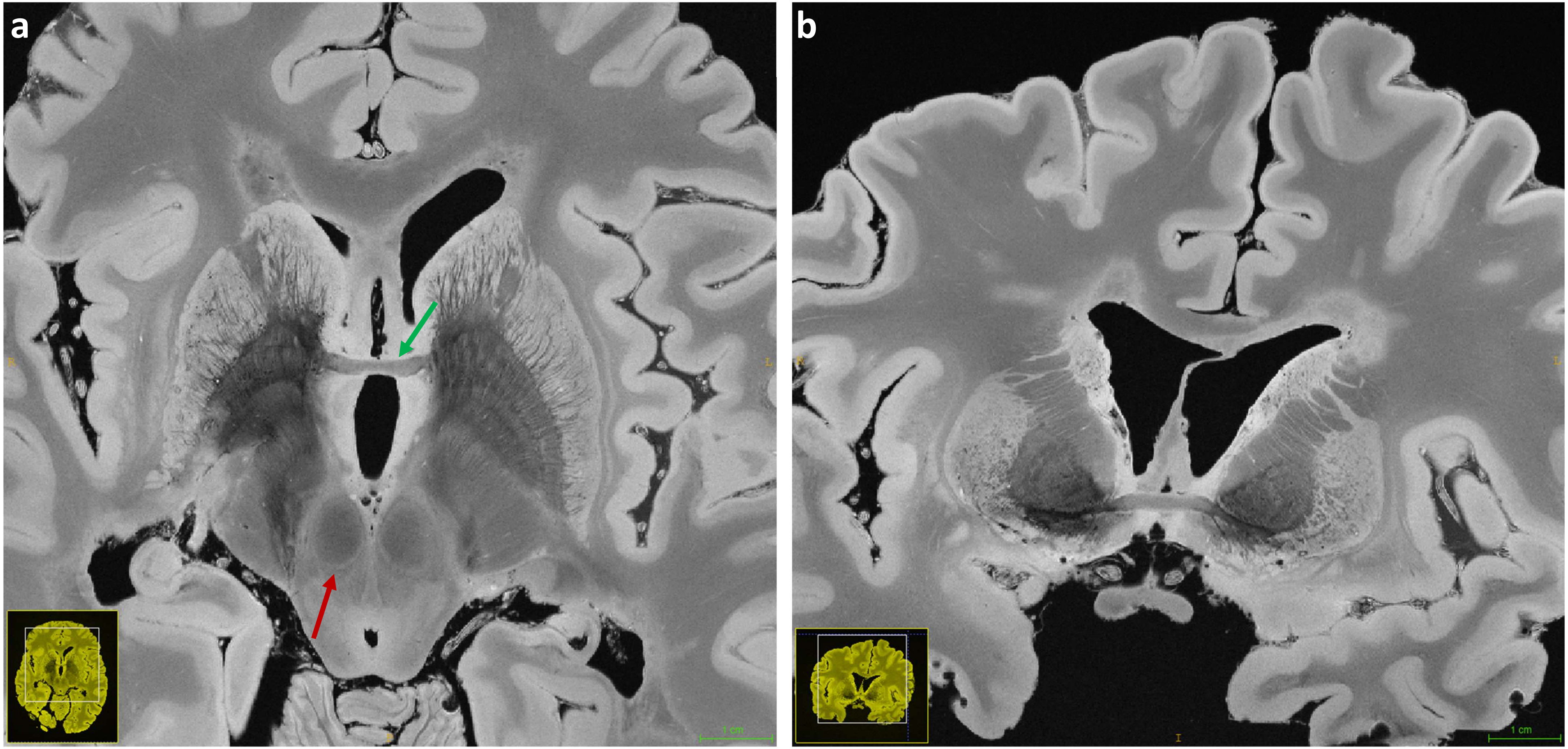


**Supporting Figure S4:** Zoomed views of a transverse section (180µm resolution, **a**) and a coronal reformation of the 240µm dataset in the plane of the anterior commissure are presented (**b**). Structures like the basal ganglia and its venous network as well as the claustrum and the anterior commissure (green arrow) are easily recognized. In the transverse plane the red nuclei are very well defined (red arrow).

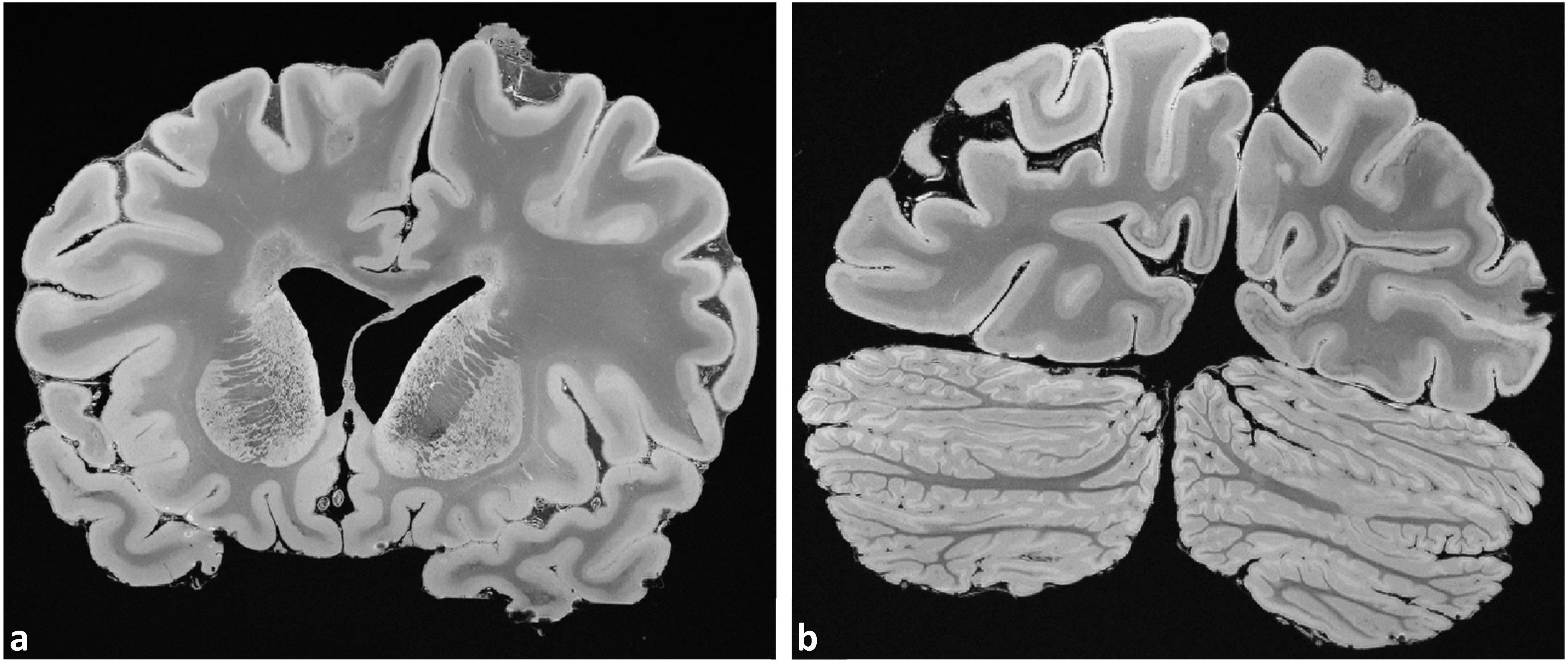


**Supporting Figure S5:** Representative coronal sections from the 200µm (**a**) and the 160µm (**b**) resolution dataset are shown. The interconnected structure of the striatum nuclei (i.e., putamen and caudate) (**a**) and the very tightly folded layers of GM of the cerebellar cortex (**b**) can be visualized in detail.
